# Propofol-induced loss of responsiveness reorganizes cortical traveling waves in the human brain

**DOI:** 10.64898/2026.04.30.721975

**Authors:** Veronica M. Zarr, Tyler S. Davis, Paul A. House, Bradley Greger, Zachary W. Davis, Elliot H. Smith

## Abstract

Consciousness depends on the coordinated propagation of neural activity across cortical networks. Traveling waves, the spatiotemporal patterns of phase-aligned neural activity, are thought to support large-scale cortical communication, yet how these dynamics are altered during general anesthesia remains poorly understood. Here, we examined the effects of propofol on cortical traveling waves recorded with high-density microelectrode arrays in the temporal lobes of two human participants (both men). We identified and characterized changes in neural traveling wave properties as participants underwent general anesthesia. Propofol produced robust, state-dependent reorganization of wave dynamics: increased propagation speed, shifted propagation directions, and altered spectral structure. At the neuronal level, propofol was accompanied by pronounced changes in firing activity and spike-wave relationships, linking mesoscale traveling waves to coordinated changes in neuronal firing dynamics. Together, these findings demonstrate how propofol reshapes the spatiotemporal organization of cortical activity across spatial scales.

## Introduction

How anesthetic drugs such as propofol induce unresponsiveness and amnesia remains one of the central unresolved questions in systems neuroscience. Although the molecular mechanisms of propofol, such as its potentiation of GABAergic inhibition, are well characterized, how such synaptic effects scale to disrupt large-scale neural dynamics, giving rise to a common anesthetic endpoint, is far less understood^1,2^. Pioneering work has shown that propofol profoundly alters cortical activity, producing large-amplitude slow oscillations, anteriorization of alpha rhythms, and hypersynchronous network states that fragment communication across the distributed cortical and subcortical systems implicated in conscious processing^3–6^. While anesthesia is known to induce fragmented states at both macroscopic and cellular scales that impede long-range communication there is a critical need to understand how these disruptions manifest across concurrent spatial scales^6–8^.

Neural traveling waves provide a powerful framework for understanding these disruptions because they reflect the emergent coordination of neuronal populations across space and time^9^. Neural traveling waves have been proposed to implement a canonical computation across spatial scales, organizing neural activity across the cortical hierarchy^10^. Through providing sub-threshold depolarization to individual neurons, traveling waves have been shown to modulate neuronal firing probability in local tissue^11,12^. In the awake state, it is thought that waves provide the spatiotemporal scaffolding necessary for the flexible routing of information across cortical networks to support sensory integration, attention, memory, and motor planning^13–19^. Because traveling waves link population-level dynamics to the timing of neuronal firing, alterations in their propagation patterns may represent a fundamental mechanism through which anesthetics reshape cortical information processing.

Despite growing evidence that traveling waves are altered by anesthesia^16,20–22^, traveling waves have not been directly compared between awake and propofol-LOR states with high spatiotemporal resolution in the human brain. Here, we test the hypothesis that propofol-induced loss of responsiveness (propofol-LOR) is accompanied by a systematic reorganization of the spatiotemporal features of traveling waves. Specifically, we examine how wave direction, propagation speed, spectral dynamics, and coordination of local firing activity differ between awake and propofol-LOR states. We show that propofol profoundly alters the cortex, disrupting mesoscale spatiotemporal dynamics. Together, these findings position traveling waves as emergent systems-level correlates of consciousness, providing a spatiotemporal framework for neural communication necessary for conscious information processing.

## Materials and Methods

### Participants

Two adult patients participated in this study after providing informed consent under a protocol approved by the University of Utah Institutional Review Board. Patient 1 was a 31-year-old man and Patient 2 was a 64-year-old man, both undergoing electrode explantation after a neuromonitoring period required for surgical treatment of their drug-resistant epilepsy. Comprehensive presurgical evaluation, including seizure semiology, scalp EEG, video EEG monitoring, high-resolution MRI, and PET imaging localized seizure onset zones to mesial temporal structures in both individuals. Our recordings employed Utah-style microelectrode arrays (UMAs), consisting of a fixed grid of penetrating microelectrodes, facilitating straightforward measurement of traveling waves^23–25^. In both patients, a 10 by 10 grid of penetrating (1 mm) microelectrodes spaced 400 micrometers apart (Blackrock Microsystems, Salt Lake City, Utah) was implanted into the middle temporal gyrus, approximately 3 cm posterior to the temporal pole. Following completion of the experimental recordings, the array was removed and each patient proceeded to standard anterior temporal lobectomy. Complete surgical procedures are described in House et al.^26^.

### Local field potentials (LFPs) and multi-unit firing activity

Anesthesia during UMA implantation and data acquisition was maintained using continuous intravenous infusions of propofol and remifentanil. After implantation, infusion rates were adjusted to maintain a bispectral index (BIS) near 50, corresponding to a lighter level of general anesthesia. Following a stable baseline period, propofol boluses were administered to deepen anesthesia, reducing BIS values to approximately 20-30 and inducing transient periods of cortical suppression. Neural activity was recorded from the UMA using a Blackrock Cerebus acquisition system at a sampling rate of 30 kHz. Signals were referenced to a subdural platinum wire located more than 2 cm from the array. The recordings captured both local field potentials (LFPs) and multiunit activity (MUA). Raw data were downsampled to 2 kHz and converted to microvolts using the Blackrock NPMK toolbox. LFP signals were high-pass filtered at 1 Hz and notch filtered at 60 Hz to remove slow drifts and line noise, respectively.

MUA was extracted directly from the raw recordings sampled at 30 kHz using an established detection procedure^27^. For each channel, the broadband signal was band-pass filtered between 300 and 3,000 Hz using a zero-phase finite impulse response filter to isolate the MUA band. The filtered signal was mean-centered, and candidate multiunit events were identified as negative deflections whose amplitudes exceeded a channel-specific threshold defined as four times the root-mean-square (RMS) of the filtered signal in the negative direction. MUA event times were detected as local minima crossing this threshold and converted to timestamps. No single-unit sorting was performed; all detected events were treated as multiunit activity and retained on a per-channel basis for subsequent analyses.

Noisy channels were removed using principal component analysis, in which channels with extreme loadings on the first principal component, representing outlying shared covariance, were removed. Visual inspection of the neural recordings confirmed the absence of ictal and interictal firing activity. Accordingly, no additional analyses to exclude epileptiform activity were performed, consistent with established protocols for using epileptic recordings to study healthy human brain function^28–30^. Electrode positions were obtained from the manufacturer’s pinout map.

### Traveling wave identification and characterization

To detect traveling waves across the array, we used generalized phase (GP),^17^ which captures the instantaneous phase of broadband LFP fluctuations (1-40 Hz). GP was computed for each microelectrode and assembled into a spatiotemporal phase field *ϕ*(*x,y,t*), where *x* and *y* denote electrode coordinates and *t* denotes time. Candidate wave events were evaluated at discrete time points (“evaluation points”) sampled from the continuous GP time series. At each evaluation point, the spatial phase field was characterized by estimating both a planar phase gradient and an effective wave source location. The source was defined as the location minimizing the squared error between observed phase values and a radially propagating phase model. Distances from this source to each electrode were then computed based on known electrode spacing.

The spatial organization of phase was quantified using a phase-distance correlation (ρ), which measures the relationship between instantaneous phase values and distance from the estimated source. A traveling wave was operationally defined as an evaluation point at which the observed phase-distance correlation exceeded that expected under a null distribution generated by randomly permuting phase values across electrode positions (500 iterations; criterion: *α* = 0.05). To prevent overcounting of temporally overlapping detections, evaluation points were clustered in time using a window equal to half the median inter-evaluation point time interval, and within each cluster only the wave with the maximum ρ was retained. Results were robust to the wave selection method (maximum ρ, random ρ, or median ρ).

All hypotheses were tested within-participant and findings were cross validated across participants, as is convention in primate neuroscience. For both participants, awake and propofol-LOR states were separated based on the timing of propofol boluses for each participant. At each evaluation point, the spatial organization of phase across the UMA was quantified by fitting a planar phase field to the observed phase values. This yielded a spatial phase-gradient vector.

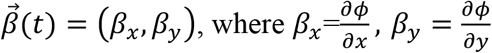

The gradient vector captures both the orientation and magnitude of phase propagation across the array. For significant waves, propagation direction was defined as the direction of decreasing phase and was quantified as the circular mean of the spatial phase gradient orientations across microelectrodes at each evaluation time. Propagation speed was computed as the ratio of the instantaneous angular frequency ω(*t*), derived from GP to the phase-gradient magnitude 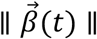.

### Statistical analyses

All statistical tests employed a criterion of *α* = 0.05. Traveling wave directions ostensibly propagated bimodally, so to characterize traveling wave direction across brain state, we fit the direction distribution for waves during awake and propofol-LOR using a two-cluster von Mises Mixture Model (vMM). The vMM is the circular analog of the Gaussian Mixture Model. Each vMM distribution can be described by a mean direction (*μ*) and a concentration parameter (*κ*), where larger *κ* values indicate tighter clustering of angles around the mean. Our vMM estimated two von Mises sub-distributions, each defined by its own *μ*, and *κ* capturing the dominant modes of wave propagation. Circular summary statistics of wave directions (e.g., circular means and resultant vectors) were computed using standard directional statistical methods via the Matlab Circular Statistics Toolbox^31^.

To characterize differences in traveling wave propagation velocity between awake and propofol-LOR states, we performed nonparametric two-sided 10,000-fold permutation testing on the median difference (propofol-LOR – awake) against a null distribution of shuffled median differences. To create the null, awake and propofol-LOR speed values were pooled, randomly shuffled, and reassigned into surrogate groups matched to the original group sizes. For each shuffle, the median difference was recomputed. The *p* value was computed as the proportion of shuffled differences whose absolute value exceeded the median difference of the true data. Waves with outlier speeds defined as larger than three times the median absolute deviation where excluded from further analysis^32^.

We used Kolmogorov–Smirnov tests to compare the overall distributions of traveling wave frequencies between brain states for each patient. We analyzed wave proportions within canonical frequency bands of δ (1-4 Hz), θ (4-8 Hz), α (8-13) and β (12-30) using two-proportion z-tests to further characterize specific frequency distribution shifts during awake and propofol-LOR.

To understand how traveling wave dynamics shaped neuronal firing, spike-phase coupling was computed by aligning neuronal firing times to the GP of the LFP for each detected wave. For each wave, MUA occurring within ± 400 ms of the traveling wave were identified, and the GP was recorded at each spike time. Spike-phases were then pooled across electrodes within that wave to obtain a population-level estimate of phase preference. Phases were binned from −π to π radians to construct per-wave spike-phase histograms, which were normalized to form probability distributions. Finally, histograms were averaged across waves within each condition to obtain the mean spike-phase distribution. To quantify preferred spike-phase, the circular mean phase was computed for each wave by taking the angle of the complex vector sum of the spike-phase histogram. Group-level preferred phase was then calculated as the circular mean across waves within each condition.

To assess differences in spike-phase coupling strength across awake and propofol-LOR states, we quantified phase modulation on a per-wave basis as the modulation depth of the spike-phase histogram, defined as the difference between the maximum and minimum probability across phase bins. Modulation depth was computed separately for each wave after pooling across directions within each condition. Waves with zero spikes were excluded. We assessed phase modulation strength across brain state by performing a two-sided 10,000-fold permutation test on the difference in medians between conditions (propofol-LOR – awake). Specifically, modulation depth values from both conditions were combined and randomly reassigned into surrogate groups of equal size, and the median difference was recomputed for each shuffle to generate a null distribution. The true median difference was then compared against this null distribution, and the *p*-value was calculated as the proportion of shuffled differences with magnitude greater than or equal to the true value. When multiple comparisons were performed, *p*-values were corrected using the Benjamini–Hochberg procedure to control for the false discovery rate. All analyses were conducted using Matlab R2023b (The MathWorks) using custom scripts adapted from the generalized-phase toolbox^17^.

## Results

We recorded from high density Utah-style microelectrode arrays in the middle temporal gyri of two adult pharmacoresistant epilepsy patients who underwent neurosurgery to treat their drug-resistant epilepsy (Fig. 1A). We measured LFP and MUA activity across all 96 microelectrodes on each array as patients became unresponsive in response to a bolus of propofol used to induce a state of general anesthesia (Fig. 1B-D). Fluctuations in GP were not synchronous across the sampled cortical area. Instead, peaks in the wideband filtered LFP (1-40 Hz) often propagated coherently across electrodes, consistent with traveling waves, during both awake and propofol-LOR (Fig. 1E). We defined a traveling wave as a nonzero spatial phase lag radiating from a putative source, identified by phase-distance correlations exceeding the 95th percentile of a permutation-derived null distribution (Fig. 1F). Across 1,991 detected evaluation points for patient 1, we identified 1,066 traveling waves after permutation testing (53.5 %), including 726 during awake and 340 during propofol-LOR. Across 1,938 detected evaluation points for patient 2, we identified 1,174 traveling waves after permutation testing (60.6 %), including 855 during awake and 319 during propofol-LOR.

**Figure 1.**
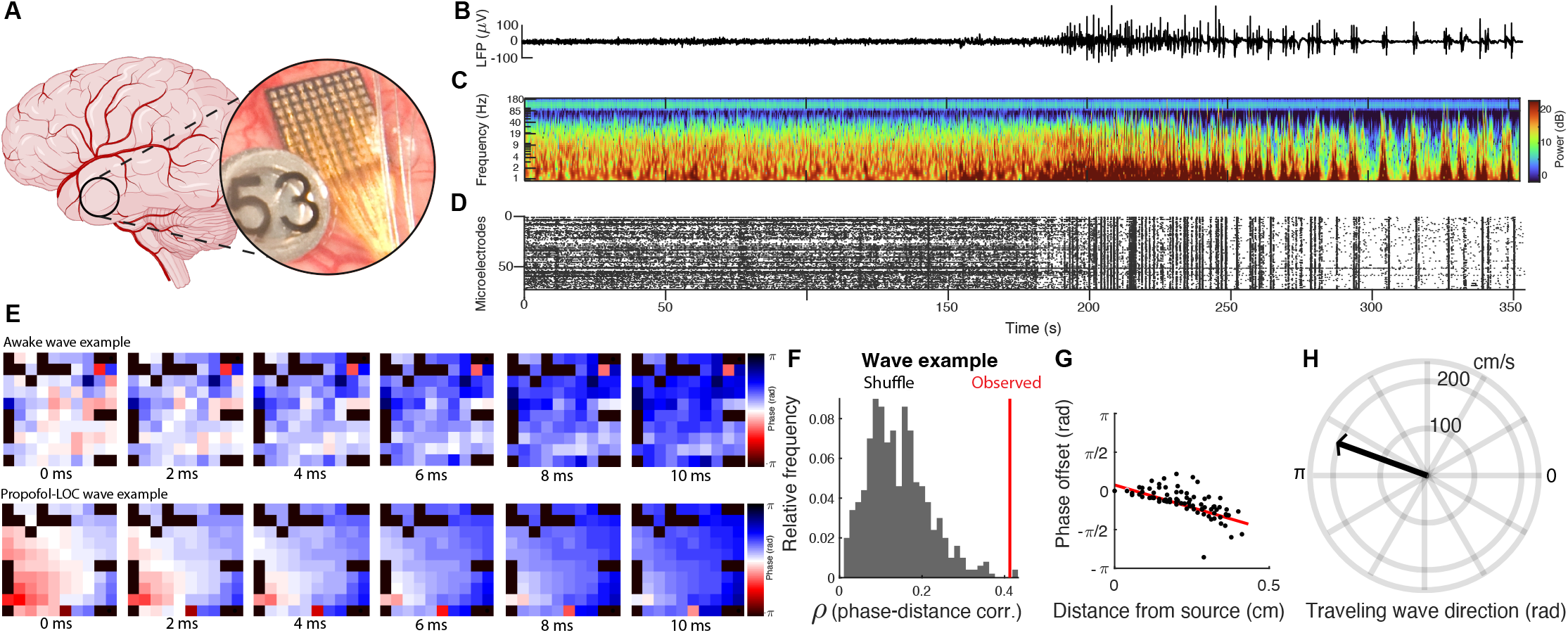
LFP fluctuations often travel as waves across the cortex across brain states. ***A***, Left: schematic of UMA placement onto the middle temporal gyrus. Right: close-up photo of the UMA (4x4 mm, 96 electrodes), implanted during a craniotomy. ***B***, Representative pre-processed UMA recording of a neurosurgical patient as he received a bolus of propofol at approximately 150 s; broadband mean LFP signal averaged across channels of the UMA. ***C***, Spectrogram of mean LFP activity across the UMA. ***D***, Raster plot of multiunit event times across the UMA. ***E***, When viewed simultaneously across the multielectrode array, spontaneous fluctuations in the wideband (1-40 Hz) LFP organize into traveling waves, visualized as instantaneous phase of the generalized-phase signal across electrodes (color scale, -*π* to *π*). Example waves from the intervals highlighted by the red and blue boxes are shown. Top, representative traveling wave during the awake state. Bottom, representative traveling wave during propofol-LOR. ***F***, Visualization of permutation test used to identify the traveling wave. Red: ρΦd of the true data. Black: ρΦd values from spatially shuffled data. ***G***, Scatter plot showing phase offsets with distance from the putative source of the detected propofol-LOR wave (black dots). The slope of the linear fit corresponds to the speed of the wave. ***H***, Quiver plot summarizing and visualizing the speed (cm/s) and direction (radians) for the example wave.

For each detected traveling wave, we characterized its spatiotemporal properties. The slope of the phase-distance relationship provided an estimate of wave speed (Fig. 1G), and wave direction was estimated using the vector gradient across the UMA (Fig. 1H). Together, these observations demonstrated that spontaneous LFP fluctuations frequently propagated as traveling waves across the cortical surface in both awake and propofol-LOR states.

Next, we characterized the features of these neural traveling waves between brain states. Applying this procedure across thousands of waves enabled us to quantify how the direction, speed and frequency of traveling waves reorganized during awake and propofol-LOR states. Traveling wave propagation direction differed between awake and propofol-LOR in both participants (Fig. 2A). Polar histograms revealed structured, non-uniform directional distributions in each state and both patients. More specifically, waves traveled antipodally along a dominant axis in both patients and brain states. To quantify these directional changes, we fit vMM models to the traveling wave direction distributions (Fig. 2B). A two-cluster vMM model captured the bimodal structure observed in both states and a direction vector denoting the circular mean of the two-mixture component means, which should be interpreted as being orthogonal to the axis of propagation.

**Figure 2.**
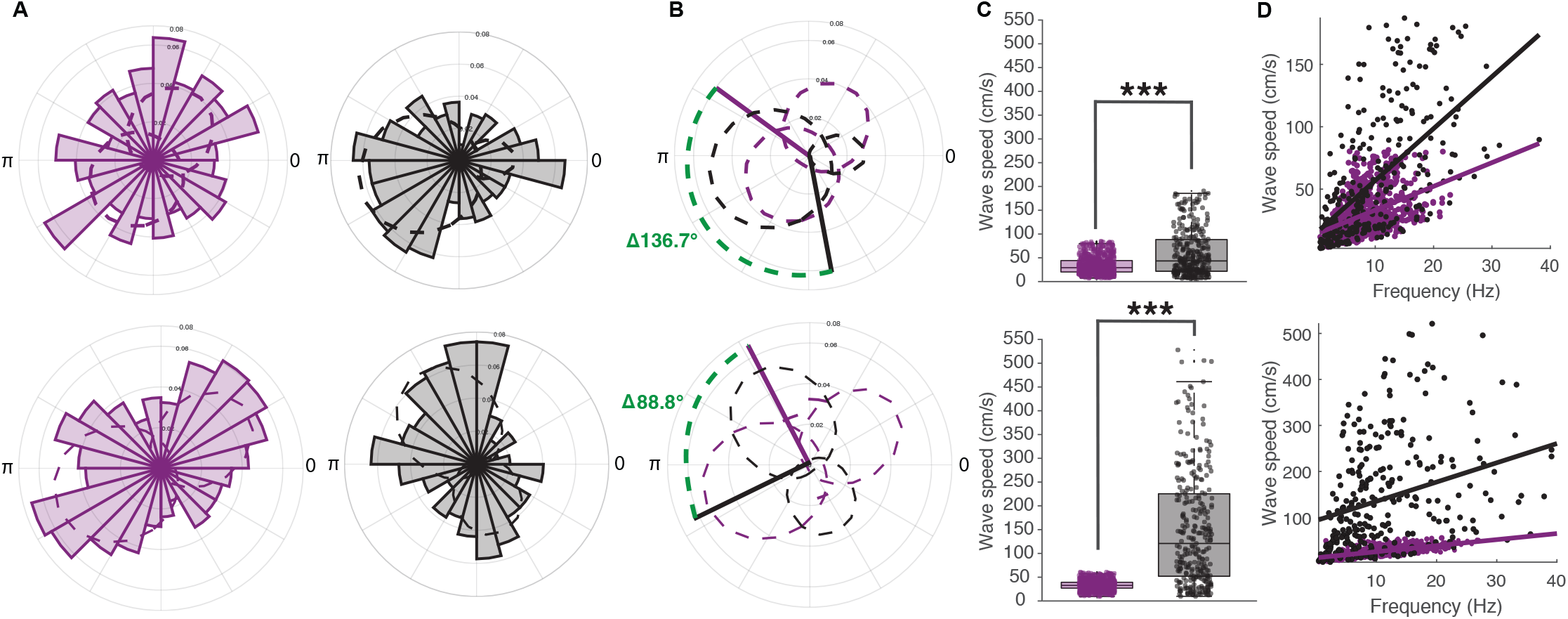
Propofol-LOR alters traveling-wave direction and speed. ***A***, Polar histograms showing the distribution of traveling wave directions during awake (purple) and propofol-LOR (black). Top, data from patient 1; bottom, data from patient 2. ***B***, Directional distributions depicted as fitted two-component von Mises mixture models (dashed lines). Solid lines represent the mean direction across waves. The difference in mean direction between states is shown in green and indicated by the dashed line connecting the awake and propofol-LOR direction vectors. ***C***, Scatter and box plots of traveling-wave propagation speeds during awake (purple) and propofol-LOR (black). Boxes denote the medians and interquartile ranges; whiskers indicate 1.5× the interquartile range. Asterisks indicate significant differences between states (\*\*\**p* < 0.001); permutation tests). ***D***, Scatter and linear regression line fit of traveling-wave propagation speeds across frequency during awake (purple) and propofol-LOR (black) for patient 1(top) and patient 2 (bottom).

During the awake state, Patient 1 exhibited a mixture mean direction of *μ* = −54.0° (−0.94 rad) and patient 2 showed a mixture mean direction of *μ* = −27.3° (−0.48 rad). Following induction of propofol-LOR, the preferred propagation axes shifted orthogonally. In patient 1, the mixture mean rotated to *μ* = 169.3° (2.95 rad), corresponding to a −136.7° (−2.34 rad) shift relative to awake (computed as the circular difference). In patient 2, the mixture mean shifted to *μ* = −116.2° (−2.03 rad), representing a −88.8° (−1.55 rad) circular shift during propofol-LOR. Across patients, the preferred propagation axis shifted by a mean of 112.8° ± 33.9°, corresponding to roughly an orthogonalization of wave directions in response to propofol-LOR. We did not observe consistent directional changes between brain states in each patient across canonical frequency bands (Supplementary Fig. 1).

Propagation speed also differed significantly between brain states (Fig. 2C). Traveling wave speeds were significantly faster during propofol-LOR in both patients (permutation tests, p < 0.001). During the awake state, Patient 1 exhibited a median wave speed of 25.75 cm/s (IQR: 16.85–40.66) and patient 2 showed a median wave speed of 25.92 cm/s (IQR: 20.27–32.43). Following induction of propofol-LOR, median speeds increased. In patient 1, the median wave speed increased to 39.95 cm/s (IQR: 18.26–84.88) corresponding to a 14.21 cm/s median wave speed increase relative to awake. In patient 2, the median wave speed increased to 114.12 cm/s (IQR: 20.27–32.43) representing a 88.20 cm/s median shift in wave speed relative to awake. We also observed significant differences in wave propagation velocity across canonical frequency bands, indicating a robust state-dependent and frequency independent modulation of propagation velocity (permutation tests, all p < 0.001; Supplementary Fig. 2).

We next asked whether the observed increase in wave speed during propofol-LOR could be explained by changes in the frequency composition of traveling waves. We regressed wave speed across frequency (Fig. 2D) and found that in both awake and propofol-LOR, wave speed increased with frequency, however, waves during propofol-LOR exhibited consistently higher propagation speeds than awake waves at similar frequencies during the awake state. In patient 1, linear regression revealed a significant positive relationship between frequency and wave speed in the awake state (β = 1.32, *t* = 12.81, *p* < 0.001, R^2^ = 0.19), which became markedly steeper under propofol (β = 4.19, *t* = 12.56, *p* < 0.001, R^2^ = 0.34; Δβ = 2.87 between states). Similarly, in patient 2, the slope increased from β = 1.99 (*t* = 26.31, *p* < 0.001, R^2^ = 0.45) in the awake state to β = 4.16 under propofol (*t*= 6.83, *p* < 0.001, R^2^ = 0.13; Δβ = 2.17 between states). These results indicate that although wave speeds linearly increased with frequency in both states, the relationship is significantly stronger under propofol, with speed differences persisting at comparable frequencies. This suggests that the increase in propagation speed reflects propofol-induced modulation of wave dynamics rather than simply a shift in frequency content. Together, these findings demonstrate that propofol-LOR is associated with pronounced shifts in traveling wave direction and increased propagation speed across human cortex.

The overall distributions of wave frequencies differed significantly between awake and propofol-LOR in each patient (two-sample Kolmogorov–Smirnov tests; Patient 1: D = 0.167, p < 0.001; Patient 2: D = 0.148, p < 0.001; Fig. 3A). Band-wise summaries across both patients revealed shifts across multiple frequency ranges following propofol, including an increases in the proportion of δ-band waves, accompanied by reductions in α-band waves in both patients (Fig. 3B; Two-proportion z-test, p < 0.001). These results point to a global broadening of traveling wave frequencies under propofol-LOR.

**Figure 3.**
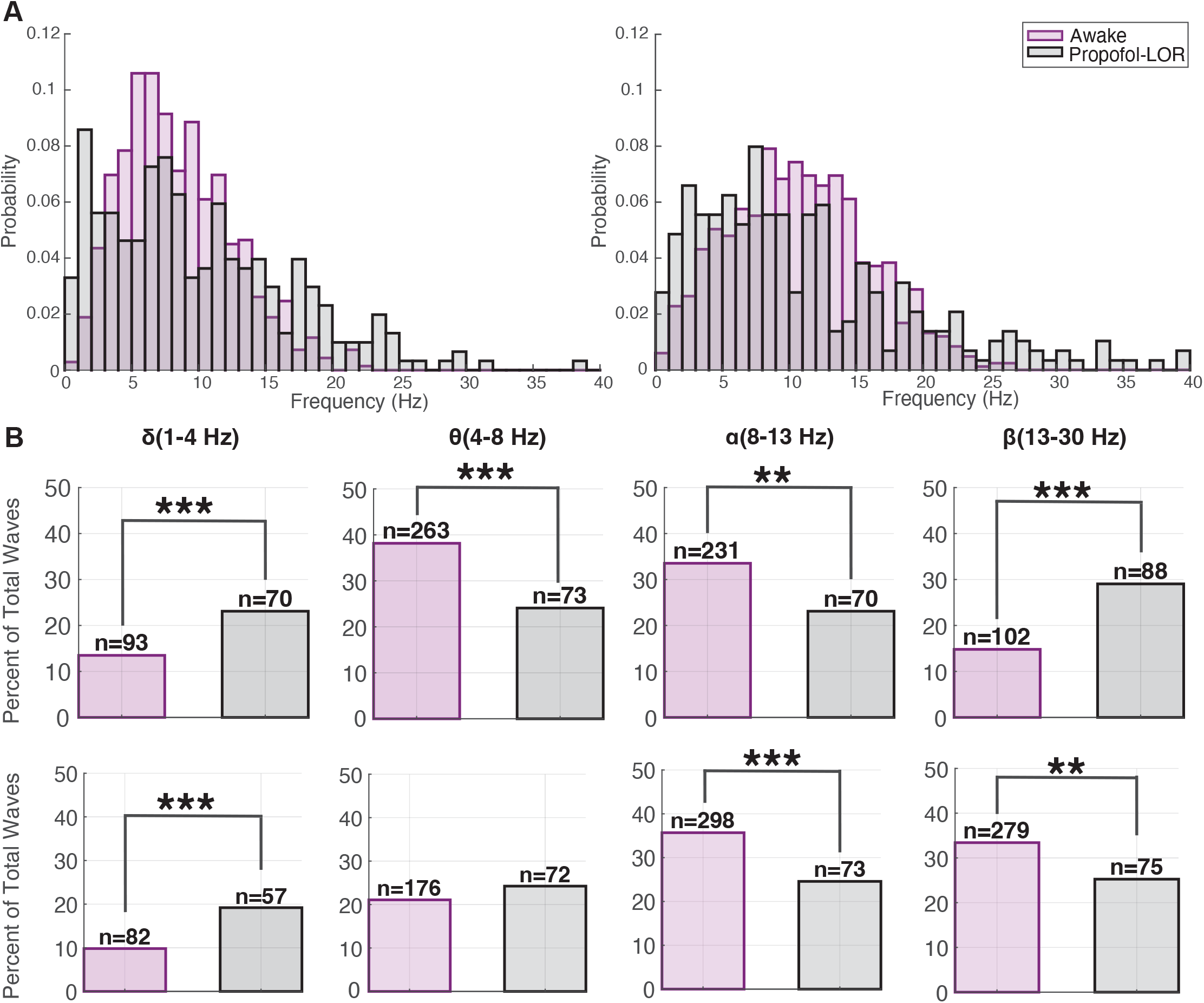
Propofol changes wave frequency distributions. ***A***, Probability density histograms of traveling wave frequencies for each patient during the awake and propofol-LOR states for patient 1 (left) and patient 2 (right). ***B***, Relative proportions of waves within canonical frequency bands (δ, θ, α, β) in the awake and propofol-LOR states for patient 1 (top) and patient 2 (bottom). Asterisks denote statistical significance (\*\**p* < 0.01, \*\*\**p* < 0.001).

Finally, we assessed how traveling waves interacted with neuronal firing between states. Circular histograms of spike-phase coupling showed that MUA in both awake and propofol-LOR conditions preferentially occurred near ± *π* radians of the generalized phase (patient 1: awake: −2.90 rad ± 0.50 rad, propofol-LOR: 2.93 rad ± 0.70; patient 2: awake: −2.90 rad ± 1.12 rad, propofol-LOR: 2.99 rad ± 1.00 rad), corresponding to a consistent preference for cells to fire at the trough of local traveling waves (Fig. 4A). Spike-phase coupling, however, was significantly stronger during propofol-LOR compared to the awake state (two-sided permutation tests, p < 0.001; Fig. 4B), indicating enhanced phase locking of local firing activity to traveling waves under anesthesia and across all frequency bands (Supplementary Fig. 3). Together, these findings demonstrate that propofol-LOR is associated not only with altered spatiotemporal wave dynamics but also with mesoscale modulation of local firing activity in concert with each traveling wave.

**Figure 4.**
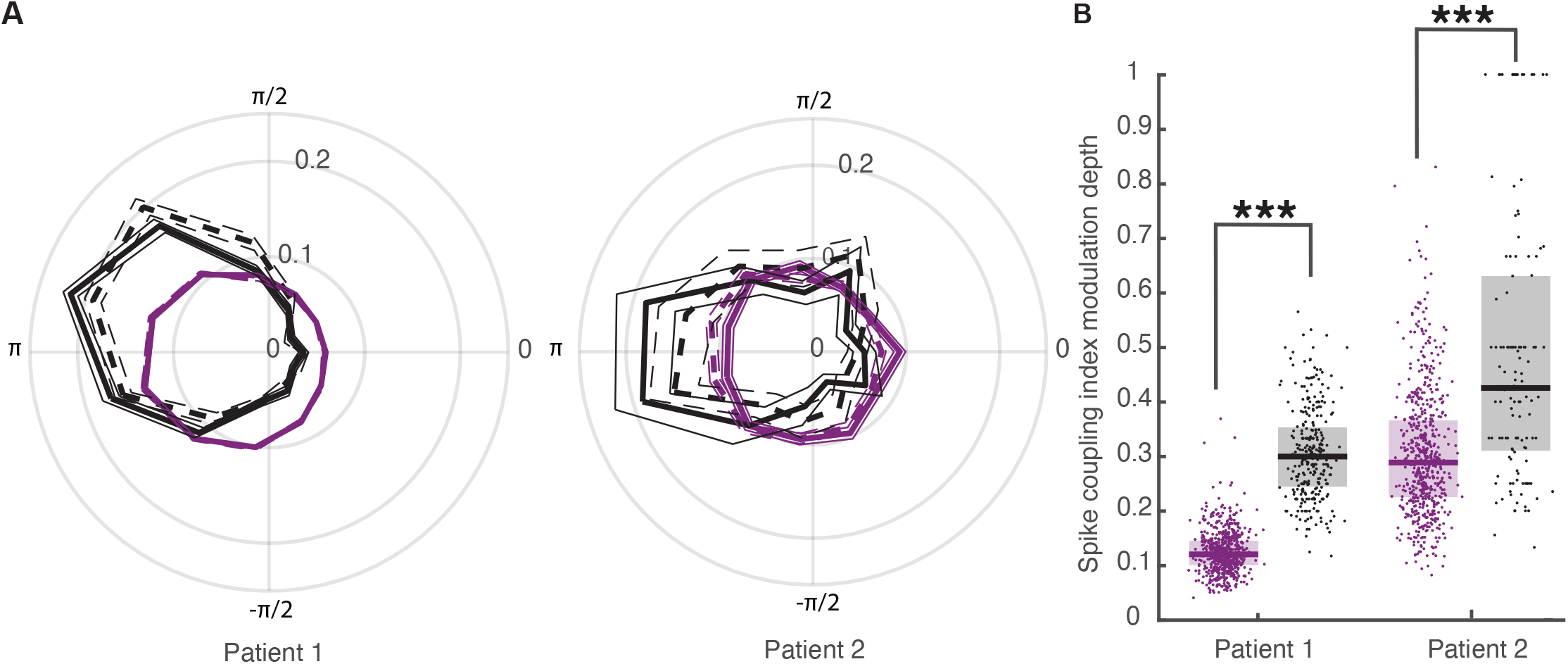
Spike-phase modulation increases during propofol-induced loss of responsiveness. ***A***, Mean spike-phase histograms showing the relative frequency of spikes as a function of generalized phase in the awake (purple) and propofol-LOR (black) states for patient 1 (left) and patient 2 (right). Dashed vs solid lines represent spike coupling to each of the two von Mises mixture model components of wave directions. Thinner lines denote the SEM around the mean. ***B***, Scatter and box plots of spike-phase coupling across all waves in patient 1 (left) and patient 2 (right) during propofol-LOR (black) compared to the awake state (purple). Boxes denote the medians and interquartile ranges. Asterisks denote statistical significance (\*\*\**p* < 0.001; permutation tests).

## Discussion

The results presented here demonstrate that propofol-LOR causes systematic reorganization of the spatiotemporal dynamics of human cortex. As patients underwent general anesthesia, traveling waves over a small patch of human association cortex orthogonalized, accelerated, broadened in frequency, and exhibited stronger coupling with action potentials. These findings thus link the anesthetic induced loss of information at the neuronal level to larger scale mechanisms of cortical information flow.

As in numerous other human traveling wave studies, we observed bimodally propagating traveling waves^16,18,19,22,23,33^. We suggest that these bimodal distributions are more likely to reflect shifting axes of large-scale activity gradients, rather than changes in local sources, as both sub-distributions of traveling waves shifted together in response to propofol^10,34^. These results suggest, that despite their advantageous high density grid-like organization, UMAs may be too small to capture the appropriate spatial scale of cortical traveling waves. In contrast to non-human primate findings, which report the emergence of bimodal traveling wave distributions only after propofol-induced loss of consciousness, our human data reveal a persistent bimodal directional organization present in both the awake and propofol-LOR states, suggesting potential species or location differences^35^. Directional biases in traveling waves have previously been reported during anesthesia and sleep and may reflect dominance of specific anatomical or functional pathways when cortical dynamics become less diverse and more stereotyped^3,12,21,22^.

Such shifting traveling wave directions under propofol were associated with wave speed increases across all frequency bands. Similar increases in wave speed have been reported during slow oscillations in sleep and anesthesia and are thought to reflect more homogeneous recruitment of neuronal populations^22,36,37^. This relationship may reflect a positive feedback loop, where increased activity in local neuronal populations around the trough of traveling waves may strengthen spatiotemporal gradients in the brain, in turn accelerating traveling waves. In the current study, wave velocities ranged from 1–530 cm/s, consistent with previously reported mesoscopic traveling waves reflecting expected conduction speeds of unmyelinated and myelinated cortical fibers^9,38^. Similar velocity ranges have been observed in human microelectrode recordings of interictal discharges^33^ and spontaneous neocortical alpha and theta oscillations^14^. The observation that propofol-LOR increases wave speed across all frequencies extends prior work showing that higher-frequency traveling waves propagate faster, and suggests that propofol shifts wave dynamics beyond frequency-dependent scaling^14,39^.

Propofol anesthesia altered the frequency distribution of traveling waves, characterized by an increase in lower-frequency delta (1-4 Hz) waves and a decrease in higher-frequency alpha (8-13 Hz) waves. Our findings are consistent with prior studies on the effects of propofol and the properties of traveling waves. The large increase in lower-frequency waves during propofol-LOR is consistent with prior reports of anesthesia related spectral power and coherence shifts and recapitulates non-human primate wave analyses with propofol-anesthesia. The reduction in alpha-frequency waves is also in line with previous studies showing that propofol-LOR is associated with a suppression of resting-state alpha rhythms^22,40^ which may reflect a disruption of feedback processes thought to support conscious awareness^15,16,41^.

Our data establish a direct link between human mesoscale traveling waves and single-unit firing activity. Under surgical anesthesia, neurons are forced into alternating “on” and “off” states^6^, permitting spikes only during narrow windows of a brain wave’s excitable phase. This increasingly strict phase locking diminishes the diverse repertoire of microstates that neuronal ensembles can produce^8^. This firing entrainment represents a mechanism by which anesthesia severely reduces the coding capacity of the human cortex^42^. Whether action potential sequences across neuronal ensembles are also disrupted by anesthesia is likely, based on changes in wave direction, speed, and spike coupling reported here, yet remains to be demonstrated empirically.

While spectral, spike-coupling and spatiotemporal features of neural traveling waves reported here were reproduced between participants, there is an important limitation. Here, we focused on propofol’s actions from spatially limited recordings in a single brain area: the middle temporal gyrus. While our recordings were spatially localized, the effects observed are likely relevant to other cortical and subcortical regions given the widespread network-level changes induced by general anesthetics.

The mechanisms implicated here may generalize to other anesthetics that primarily enhance GABAergic inhibition, including sevoflurane, isoflurane, desflurane, barbiturates, and etomidate^43,44^. In contrast, anesthetics with distinct molecular targets, such as NMDA antagonists including ketamine and nitrous oxide, produce different large-scale dynamics and may engage partially distinct mechanisms^45,46^. Future work will be necessary to directly test this hypothesis by applying the same mesoscale traveling-wave analyses across different anesthetics within comparable experimental frameworks. Such studies will be critical for determining whether changes in traveling-wave dynamics reflect a shared network-level signature of anesthetic-induced unresponsiveness or whether distinct anesthetics produce dissociable mesoscale signatures despite a common anesthetic endpoint.

## Conclusions

Overall, our analyses reveal that propofol anesthesia fundamentally reshapes the spatiotemporal architecture of cortical activity at the level of traveling waves. These findings position traveling waves as informative, systems-level correlates of consciousness and highlight their potential utility for bridging cellular, network, and behavioral accounts of anesthetic-induced loss of responsiveness.

## Acknowledgements

We gratefully acknowledge support from the Department of Neurosurgery at the University of Utah. We thank the labs of Elliot Smith, Ben Shofty, Cory Inman, and Shervin Rahimpour for feedback and discussion. This work was supported by the National Institute of Neurological Disorders and Stroke (5T32NS115723-04); the National Institutes of Health (EY014800) and Unrestricted Grant from Research to Prevent Blindness, New York, NY, to the Department of Ophthalmology & Visual Sciences, University of Utah.

## Author contributions

E.H.S., conceived and supervised the project. E.H.S., and V.M.Z., designed the experiments, critically reviewed the methods. E.H.S., V.M.Z., and Z.W.D., developed the overall research strategy. T.D., B.G., and P.A.H., collected the Utah array electrophysiological data. T.S.D., V.M.Z., and E.H.S., contributed to data preprocessing pipelines and software development. All authors contributed to data interpretation and provided critical revisions to the manuscript.

## Notes

### Competing Interest Statement

The authors have declared no competing interest.

