## Supplementary material for "Propofol-induced loss of responsiveness reorganizes cortical traveling waves in the human brain": Supplementary_Material_FINAL.pdf

### Supplementary Materials

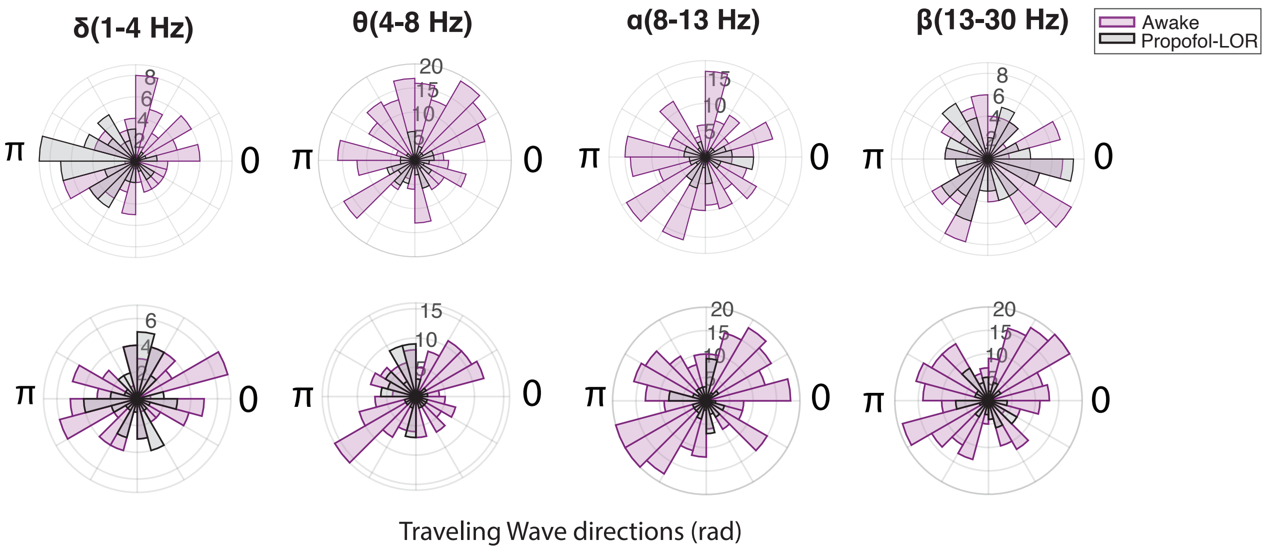


**Supplementary Figure 1. Traveling wave directions across frequency.** Polar histograms showing the distribution of traveling wave directions (rad) and wave counts stratified by frequency band during awake (purple) and propofol-LOR (black) states for patient 1 (top) and patient 2 (bottom).

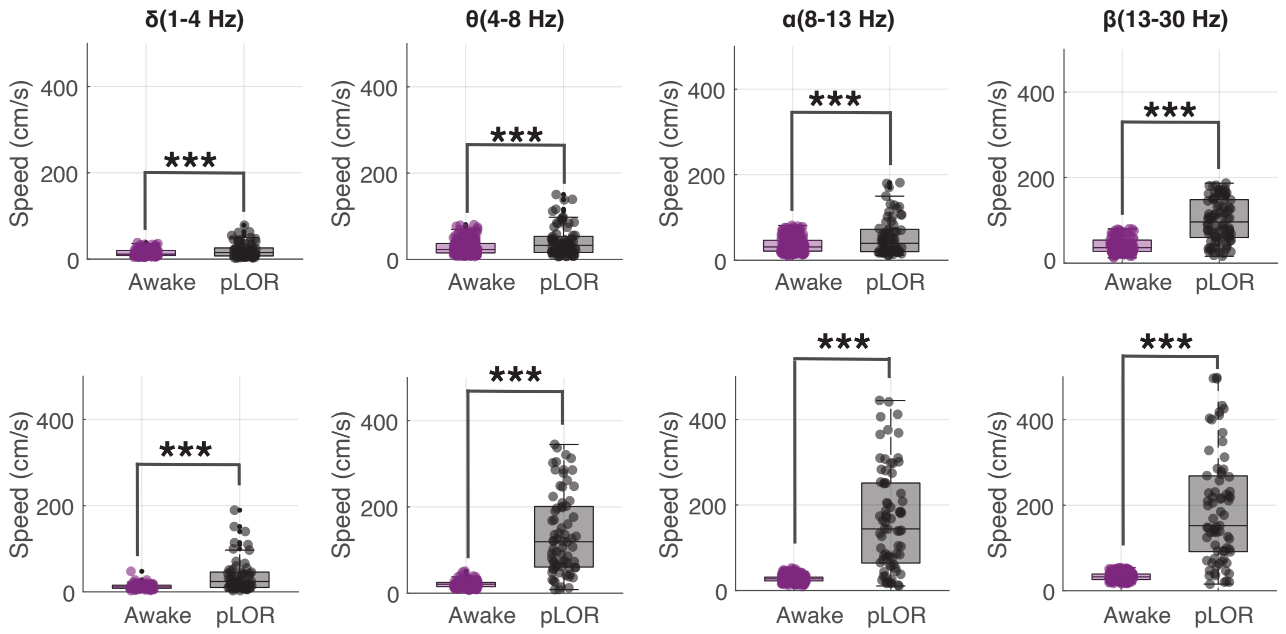


**Supplementary Figure 2. Traveling wave speeds across frequency.** Traveling wave propagation speeds during awake (purple) and propofol-LOR (black) across frequencies for patient 1 (top) and patient 2 (bottom). Asterisks denote statistical significance (***$p < 0.001$, permutation tests).

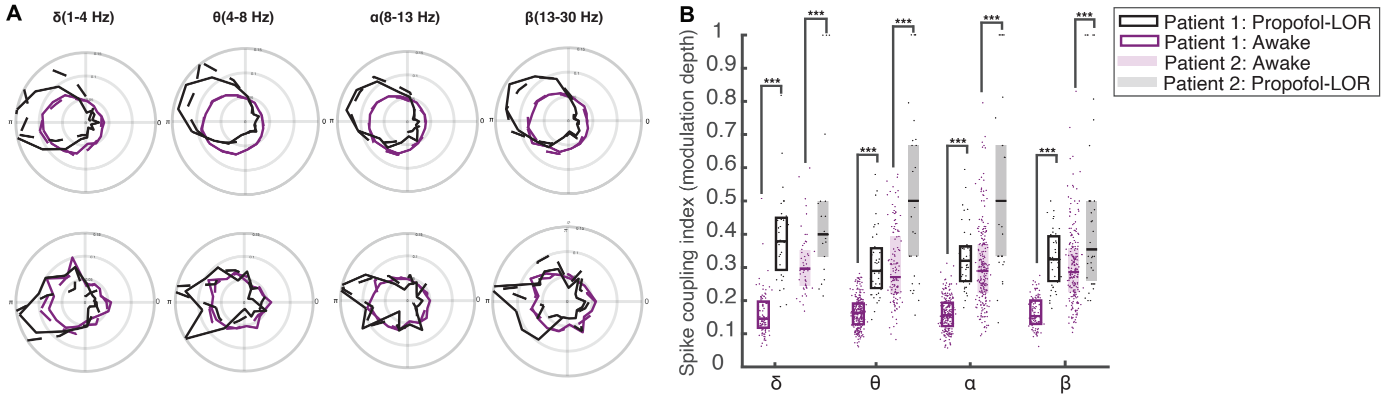


**Supplementary Figure 3. Traveling wave spike-phase modulation across frequency. (A)** Mean spike-phase histograms showing the relative frequency of spikes as a function of generalized phase in the awake (purple) and propofol-LOR (black) states, across all frequency bands for patient 1 (top) and patient 2 (bottom). **(B)** Summary of spike-phase coupling strength quantified per wave as the modulation depth of the spike-phase histogram. Coupling strength was significantly higher during propofol-LOR compared to the awake state (two-sided permutation test on the median difference, $p < 0.001$. Asterisks denote statistical significance (***$p < 0.001$).
